# Glycan-induced structural activation softens the human papillomavirus capsid for entry through reduction of intercapsomere flexibility

**DOI:** 10.1101/2024.02.01.577804

**Authors:** Yuzhen Feng, Dominik van Bodegraven, Alan Kádek, Ignacio L.B. Munguira, Laura Soria-Martinez, Sarah Nentwich, Daniel Kavan, Charlotte Uetrecht, Mario Schelhaas, Wouter H. Roos

**Affiliations:** Moleculaire Biofysica, Zernike Instituut, Rijksuniversiteit Groningen, Groningen, Netherlands; Institute of Cellular Virology, ZMBE, University of Münster, Münster 48149, Germany; CSSB Centre for Structural Systems Biology, Deutsches Elektronen-Synchrotron DESY & Leibniz Institute of Virology (LIV), Notkestraße 85, 22607 Hamburg, Germany; Institute of Microbiology of the Czech Academy of Sciences, Videnska 1083, 14220 Prague, Czech Republic; Faculty V: School of Life Sciences, University of Siegen, Am Eichenhang 50, 57076 Siegen, Germany

**Keywords:** HPV, virus entry, receptor-switching, HDX mass spectrometry, atomic force microscopy, biomechanics

## Abstract

High-risk human papillomaviruses (HPVs) cause various cancers. While type-specific prophylactic vaccines are available, additional anti-viral strategies are highly desirable. Initial HPV cell entry involves receptor-switching induced by structural capsid modifications. These modifications are initiated by interactions with cellular heparan sulphates (HS), however, their molecular nature and functional consequences remain elusive. Combining virological assays with hydrogen/deuterium exchange mass spectrometry, and atomic force microscopy, we investigated the effect of capsid-HS binding and structural activation. We show how HS-induced structural activation requires a minimal HS-chain length and simultaneous engagement of several binding sites by a single HS molecule. This engagement introduces a pincer-like force that stabilizes the capsid in a conformation with extended capsomer linkers. It results in capsid enlargement and softening, thereby facilitating L1 proteolytic cleavage and subsequent L2-externalization, as needed for cell entry. Our data will help further devising prophylactic strategies against HPV infections.

## Introduction

Human papillomaviruses (HPVs) are a large family of non-enveloped DNA viruses that infect squamous epithelia of skin or mucosa. HPV diseases range from asymptomatic infections to anogenital or oropharyngeal cancers, the latter of which are caused by so-called high-risk HPV types^1, 2, 3^. With estimated 630,000 new annual HPV-related cancer cases, they have a huge impact on public health^4^. While type-specific prophylactic vaccines have been deployed to prevent HPV infection^5^, additional strategies to fight HPV infections or related cancers remain in demand.

The life cycle of HPVs is closely linked to the differentiation of keratinocytes^6^, where entry occurs in basal cells followed by amplification, replication and transformation in spinous cells, and attenuation of genome amplification and initiation of virus assembly in granular cells^7^. The icosahedral (*T* = 7d) HPV capsids are assembled by the major and minor capsid proteins L1 and L2, respectively^8, 9, 10^. L1 forms homo-pentameric capsomers, of which 72 self-assemble into HPV particles^8, 11^. Up to 72 copies of L2 are located mostly capsid-lumenal with a few surface-exposed peptides^12, 13, 14^. Capsid stability relies on intercalation of L1 C-terminal arms of neighbouring capsomers. This link between capsomers is further stabilized by interchain disulfide bonds^15, 16, 17, 18, 19^. Due to the complex life cycle, culturing HPVs *in vitro* is similarly complex. As models for entry studies, virus-like-particles (VLPs) and so-called pseudoviruses (PsV) have been established. VLPs are formed by self-assembly of L1 into empty particles and PsVs are formed by L1/L2 VLPs encapsidating a chromatinised, reporter-gene expressing the pseudogenome^20, 21^.

HPVs bind to cells via the heparan sulphate (HS) moiety of heparan sulphate proteoglycans (HSPGs)^22, 23^. Sequential engagement by HS of distinct binding sites located on the top rim of the L1-capsomer and in the cleft between two capsomers has been suggested^24, 25^. Binding to HS but not to other highly sulphated glycosaminoglycans (GAGs) triggers an ill-defined conformational change in L1 termed “structural activation” that facilitates L1 cleavage by a secreted protease, most likely kallikrein-8 (KLK8)^26, 27^. L1 cleavage is followed by cyclophilin-assisted externalization of the L2 N-terminus from the capsid lumen^28, 29^. Proteolytic cleavage of the L2 N-terminus by cellular furin reduces capsid affinity to HSPGs, likely followed by transfer to a secondary receptor^30, 31, 32^. Integrin α6, annexin A2 heterotetramer, growth factor receptors, and the tetraspanins CD63 and CD151 have been suggested as secondary receptor candidates^33^. The particles are internalized by a potentially novel endocytic mechanism^34, 35^, and routed via the endosomal pathway and retrograde transport to the Golgi apparatus^34, 36, 37^. Finally, the DNA is delivered to the nucleus after nuclear envelope breakdown during mitosis^38, 39^.

Capsid protein reorganisation as proposed for HPVs is common in the life cycles of various viruses^15, 40, 41^. However, the nature of these changes in HPVs with regards to structural alterations and their biochemical and mechanical consequences are so far unexplored. To address these complex issues at multiple levels, we combined virological approaches, atomic force microscopy (AFM), and hydrogen/deuterium exchange mass spectrometry (HDX-MS). While AFM nanoindentation enables the measurement of viral mechanical properties at a single-particle level^42^, local changes of structure and dynamics in proteins can instead be mapped by HDX-MS ^43, 44^. Our results indicate that successful virion activation by HS binding requires a certain minimal glycan length. The glycan likely engages the virus at multiple binding sites and softens the virus particle. Softening requires engagement of HS binding sites in the cleft between virus capsomers and is reversible. Combined with our HDX-MS data showing decreased flexibility of both the N- and C-terminal arms of L1 connecting capsomers, we propose the first molecular model of HPV structural activation. This model involves a pincer-like force exerted by heparin that locks the particle in the most extended of several alternating conformations of the invading C-terminal arm, which results in enlarging and softening of the HPV capsid.

## Results

### Structural activation of HPV16 depends on glycan length

To address glycan requirements for structural activation, we made use of a facile seedover assay (Figure 1A) ^26^. Here, HPV16 PsV bound to the extracellular matrix (ECM)-resident laminin-332 are unable to infect cells with undersulphated HS moieties, e.g. by NaClO_3_-treatment^26, 45, 46^. However, if the virus is structurally activated by engagement of HS or heparin, a fully sulphated HS analogue often used as HS model, those cells are infected^26^. While short heparin oligosaccharides are able to bind to HPV16 capsids^26, 47^, it is unclear whether they structurally activate the virus as observed for long polysaccharides. Thus, we initially tested, if GAG length may be of importance for activation using heparin and HS of various lengths. While preincubation of HPV16 PsV with long heparin chains dose-dependently restored infection of NaClO_3_-treated HaCaT cells as expected (Figure 1B), short heparin oligosaccharides that had a degree of polymerization (dp) of five saccharides (Fondaparinux) or up to dp 20 (Figure 1C & D) failed to activate. As short heparin oligosaccharides engage the virus^26, 47^, the results indicate that several HS binding sites on the capsid have to be simultaneously engaged by the same GAG to structurally activate the virus for infection. As size-defined heparin oligomers longer than dp 20 are unavailable, we further probed entry using fractionated HS. Here, we used HS with a low degree of sulphation and an average dp 190 (dp190 lowS) as well as shorter HS with a high degree of sulphation and an average dp 40 (dp40 highS). HS dp190 lowS failed to activate the virus for infection (Figure 1E). On the contrary, HS dp40 highS displayed a partial activation of infection (Figure 1E) in line with partial structural activation. It is important to note that the reduced degree in activation of the HS dp40 highS fraction as compared to e.g. heparin is likely the result of a lower degree of sulphation, and perhaps the inherent length dispersity of such fractions. Cumulatively, our results indicate that multivalent engagement of several capsid binding sites by one sulphated HS molecule of a length > dp 20 promotes structural activation.

**Figure 1:**
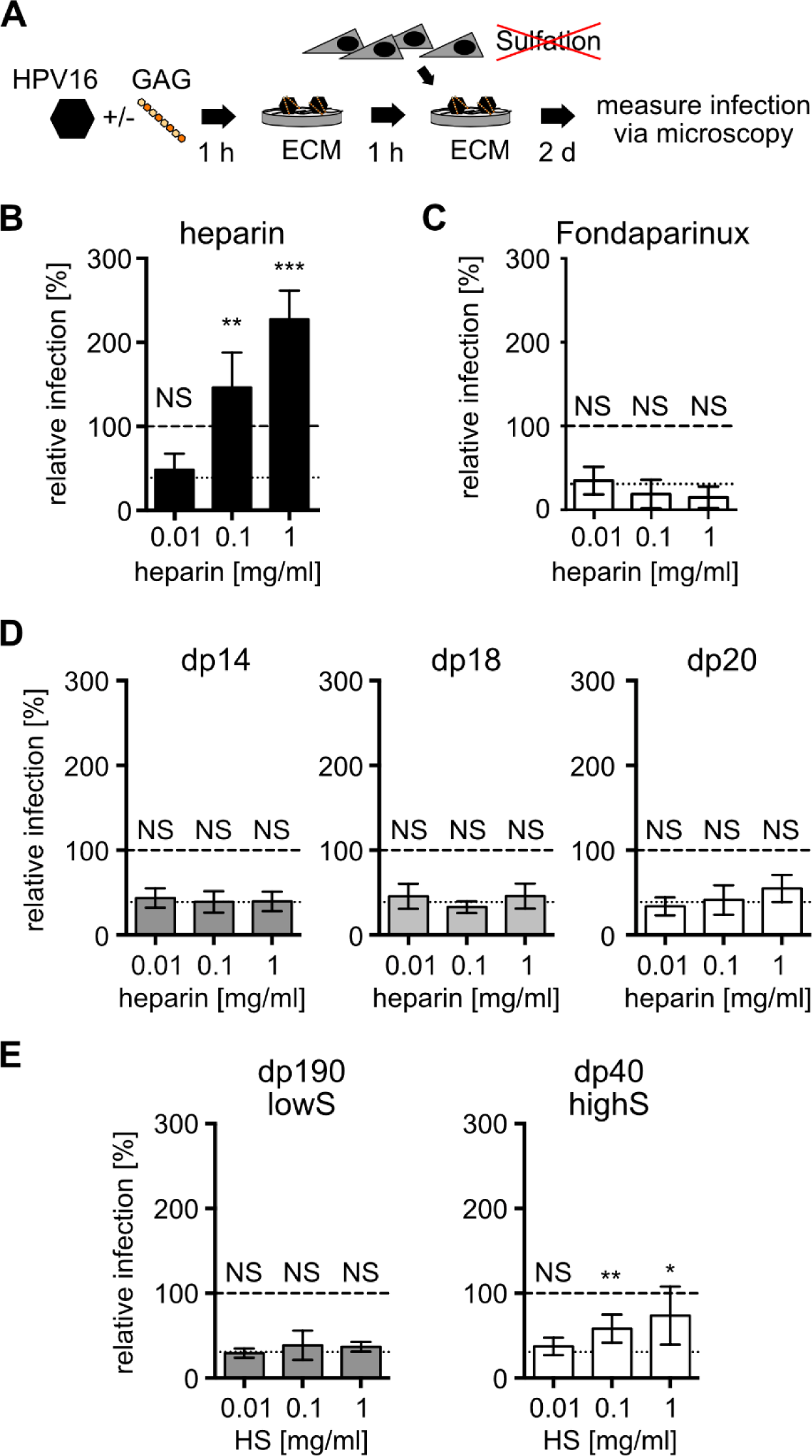
Structural activation of HPV16 is dependent on GAG length. (A) Seedover experimental schematic. Before binding to ECM HPV16 PsVs were incubated with the indicated GAG for 1 h. NaClO_3_ treated HaCaT cells were seeded on top. Infection was scored by microscopy and displayed as average of GFP-positive cells in % of total cells relative to the untreated control for three independent experiments ± standard deviation (SD). Seedover was conducted with (B) heparin, (C) Fondaparinux, (D) short heparin oligomers or (E) heparan sulfate fractions. In (E) the average of six independent experiments is displayed ± standard deviation (SD). The long dashes indicate the infection of HPV16 PsVs in untreated HaCaT cells and the short dashes in NaClO_3_ treated HaCaT cells. P values are indicated by asterisks: p<0.001 (***), p<0.01 (**), p<0.05 (*), nonsignificant (NS).

### GAGs with longer saccharide chains trigger changes in mechanical properties of HPV16

Next, we asked whether multivalent virion engagement by HS would elicit mechanical consequences in the capsid using AFM (Figure 2A). For this, we used L1-only VLPs, which engage HS/heparin as PsV^47^, but as they lack DNA, any detectable changes in mechanical properties are solely due to changes in L1 conformation upon HS engagement. In our AFM imaging and nanoindentation studies using heparin or Fondaparinux as poly- or pentasaccharide, respectively, (Figure 2B-C), HPV16 VLPs retained their overall morphology irrespective of the presence or absence of glycans. For all conditions, HPV capsomers were clearly detected by AFM (Figure 2C). Through indentation of HPV16 VLP by the AFM tip, a force-displacement (F-D) curve was recorded demonstrating elastic deformation of particles under applied force until buckling or other non-linear deformations occurred (Figure 2B). The slopes of the elastic deformation phases and the positions where non-linear deformations started, the critical force, already indicated a qualitative difference of the mechanical properties between heparin and Fondaparinux-treated particles. Quantitative assessment showed that heparin significantly decreased viral spring constant and critical force of HPV16 VLP whereas Fondaparinux did not (Figure 2D-E). Moreover, incubation with heparin significantly increased the size of HPV16 VLPs (Figure S1A). Hence, mere binding of the longer glycan triggered changes in the capsid physical properties of HPV16, which correlates with its ability to structurally activate the particle.

**Figure 2:**
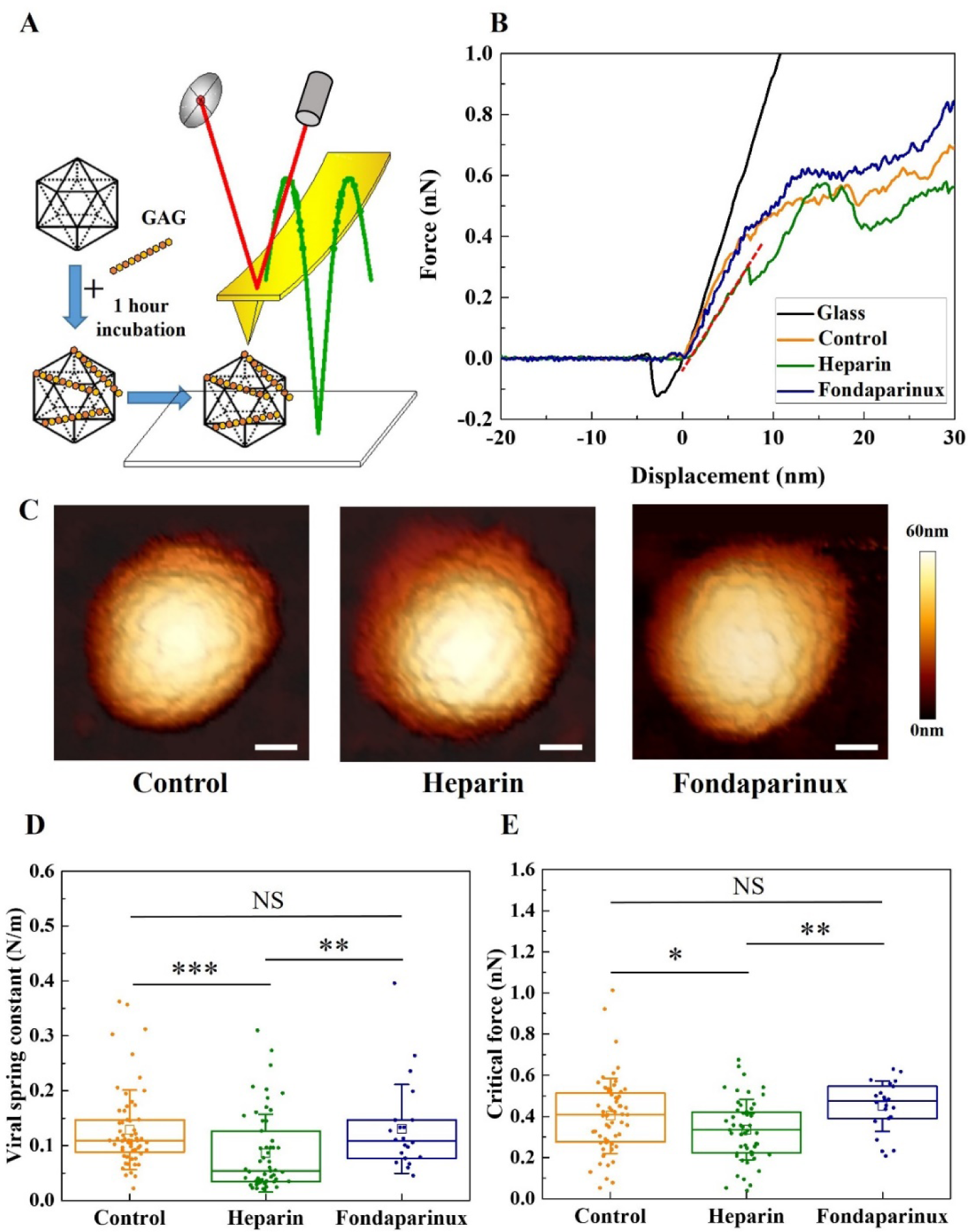
GAGs with certain lengths induce softening and enlargement of HPV capsids. (A) Schematic of the AFM measurements setup. HPV16 VLPs were incubated with GAGs for 1 hour and then imaged and indented by AFM in liquid. (B) Typical force-displacement curves acquired by using an AFM tip to indent a single VLP treated by the indicated GAG. The dark red dashed line marks the linear phase on the approach line of one of the force-displacement curves used for linear fitting. (C) Typical AFM images of the VLPs incubated with the indicated GAG before indentation. The scale bar is 20 nm. (D) Box plot of the viral spring constant and (E) box plot of the critical force of the VLPs treated by the indicated GAG. The square in the box plots indicates the mean, the top and bottom of the box are the 25th and 75th percentiles respectively, and the whiskers represent the standard deviation. The line in the box indicates the median. P values were determined for viral spring constant by Kruskal-Wallis test and for critical force by one-way ANOVA test. P values for the comparison of viral spring constant between control (N = 60 data points) and heparin (N = 55 data points), control and Fondaparinux (N = 22 data points), heparin and Fondaparinux are 1.27 × 10^−4^, 0.95, and 8.91 × 10^−3^ respectively. P values for the comparison of critical force between control and heparin, control and Fondaparinux, heparin and Fondaparinux are 2.67 × 10^−2^, 0.24, and 5.57 × 10^−3^ respectively. P values are indicated by asterisks: p<0.001 (***), p<0.01 (**), p<0.05 (*), nonsignificant (NS).

Notably, the indentation-induced non-linear deformations were reversible in our nanoindentation measurements. There were no significant changes in the morphologies and sizes of HPV16 VLPs before and after indentation (Figure S2A-B), indicating a high degree of particle robustness and flexibility withstanding loading forces up to 3 nN. The F-D curves corresponding to five successive indentations of a single particle showed that while the critical force gradually decreased, the slopes of each indentation were almost overlapping (Figure S2C). This suggests that the reversibility of HPV capsids upon deformation results from its flexible and partially dynamic capsid structure where capsomers are interconnected by a network of linking peptides. In summary, we observed a softening of the HPV16 capsids after incubation with heparin as delineated by the observed decrease in viral spring constant.

### Heparin-binding induced HPV structural activation is reversible

Since structural activation and the correlated softening of virus particles required GAGs with longer saccharide chains, we wondered whether multivalent interactions of one GAG with several sites on the viral capsid would be crucial to maintain the conformational change. To investigate a potential reversibility of structural activation, HPV16 PsV were allowed to interact with immobilized heparin, subsequently eluted, and used to perform seedover experiments. Even though these particles were thus temporarily bound to heparin, no recovery of infection was observable (Figure 3A). Likely, any structural activation triggered by transient binding to heparin was not conserved after elution.

**Figure 3:**
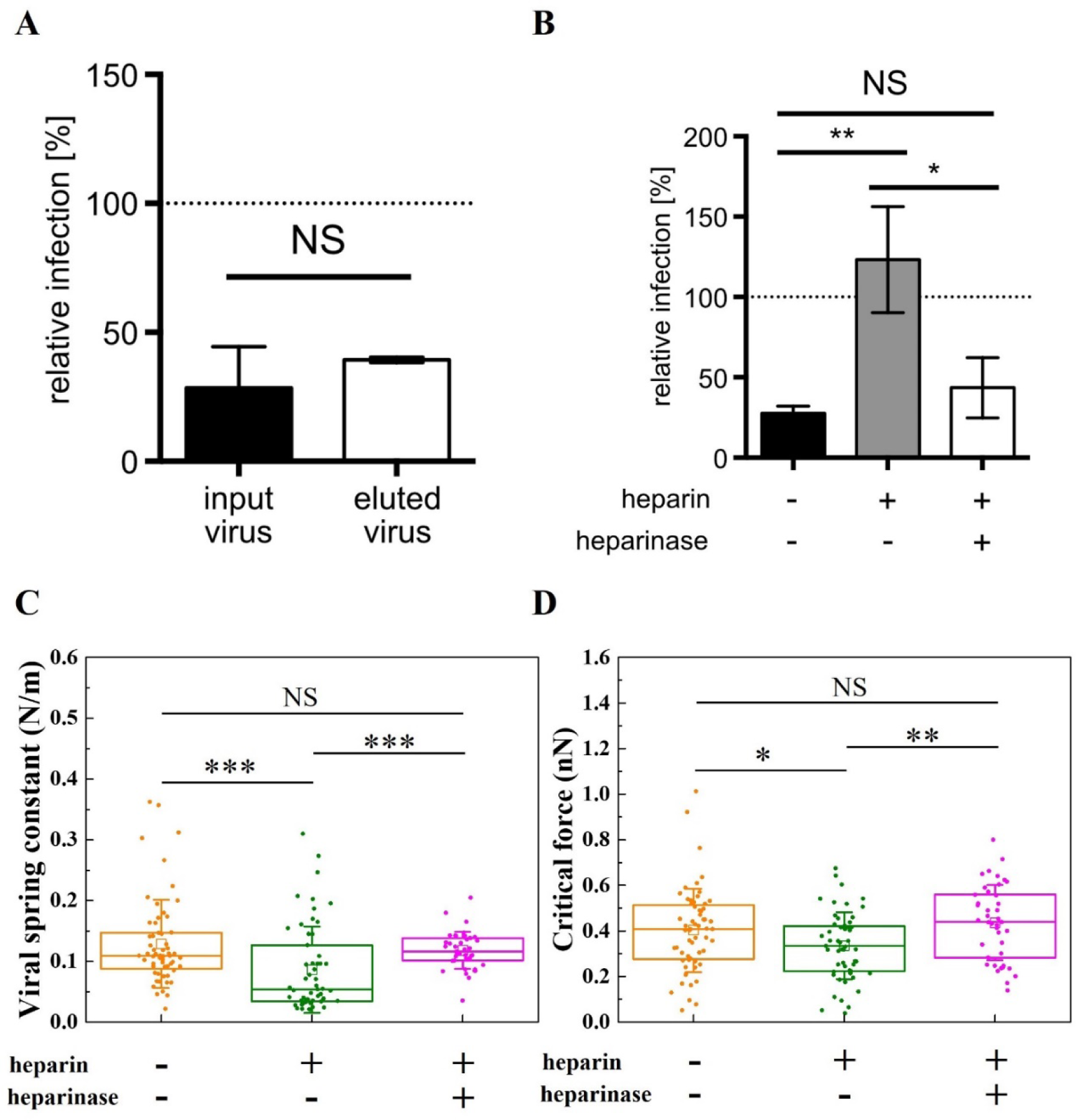
HPV structural activation induced by heparin is reversible. (A) Infection of NaClO_3_ treated HaCaT cells with PsVs before (input) or after (eluted) being subjected to heparin affinity chromatography. *N* = 3 (B) Seedover experiment results of PsVs treated with 1 mg/mL heparin and the PsVs treated with 0.5 UN heparinase subsequent to heparin treatment. *N* = 3 (C) Box plot of the viral spring constant of the VLPs treated by heparin and the VLPs treated by heparinase subsequent to heparin treatment (N = 41 data points). (D) Box plot of the critical force of the VLPs treated by heparin and the VLPs treated by heparinase subsequent to heparin treatment. P values were determined for viral spring constant by Kruskal-Wallis test and for critical force by one-way ANOVA test. P values for the comparison of viral spring constant between control and heparin, control and heparinise subsequent to heparin, heparin and heparinise subsequent to heparin are 2.76 × 10^−4^, 0.69, and 2.57 × 10^-4^ respectively. P values for the comparison of critical force between the above three pairs are 3.20 × 10^−2^, 0.32, and 3.78 × 10^−3^ respectively. P values are indicated by asterisks: p<0.001 (***), p<0.01 (**), p<0.05 (*), nonsignificant (NS).

To further corroborate this, heparin-bound HPV16 PsV were subjected to heparinase treatment cleaving heparin into oligosaccharides. In contrast to heparin-bound HPV16, heparin-unbound and heparinase-treated virus failed to exhibit structural activation as evidenced by loss of their heparin-associated recovery of infection of undersulphated HaCaT cells (Figure 3B). Moreover, a similar reversion of mechanical properties and size of HPV16 VLPs after heparinase treatment was observed by AFM-based studies. Heparinase treatment led to similar viral spring constants, critical forces and sizes as untreated HPV16, whereas heparin itself exhibited significant changes (Figure 3C-D and Figure S1B). This indicated that sustained interactions with long HS chains are indeed required to maintain the conformational change elicited by HS-virus interaction.

### K442 and K443 are essential for activation of HPV16 through heparin

So, why would multivalent binding of GAGs with longer saccharide chains be required to maintain a conformational change? Distinct HS binding sites were described in structural models from X-ray crystallography of HPV16 pentamer crystals soaked with a solution of heparin oligomers^24^. Intriguingly, these sites align and form a track that leads from the top rim into the canyon between two pentamers so that a multivalent binding of a longer polysaccharide along these sites is plausible (Figure 4A). The binding site at the top rim (lysines 278 and 361) proposed to engage HS first^25^ is furthest away from a site within the canyon between pentamers formed by lysines 442 and 443. The latter is directly connected to the invading C-terminal arm that intercalates with the adjacent pentamer. Moreover, the epitope revealed upon structural activation (recognized by the H16.B20 antibody)^26^ is located at the lower part of the invading arm. Thus, we hypothesized that a long heparin molecule needs to span several binding sites. As the binding sites lead from the top rim to the side of the pentamer, the GAG would be bent. Yet, highly sulphated HS or heparin are rather stiff molecules. The tension within heparin generated by binding to the capsid could thus generate enough force on the L1 molecule to induce a conformational change. Thus, the K442/443 site may work as a ‘handle’ for this tension-force generation as the heparin binding site directly connected to the invading arm, which is the likeliest site to result in biophysical changes if modified.

**Figure 4:**
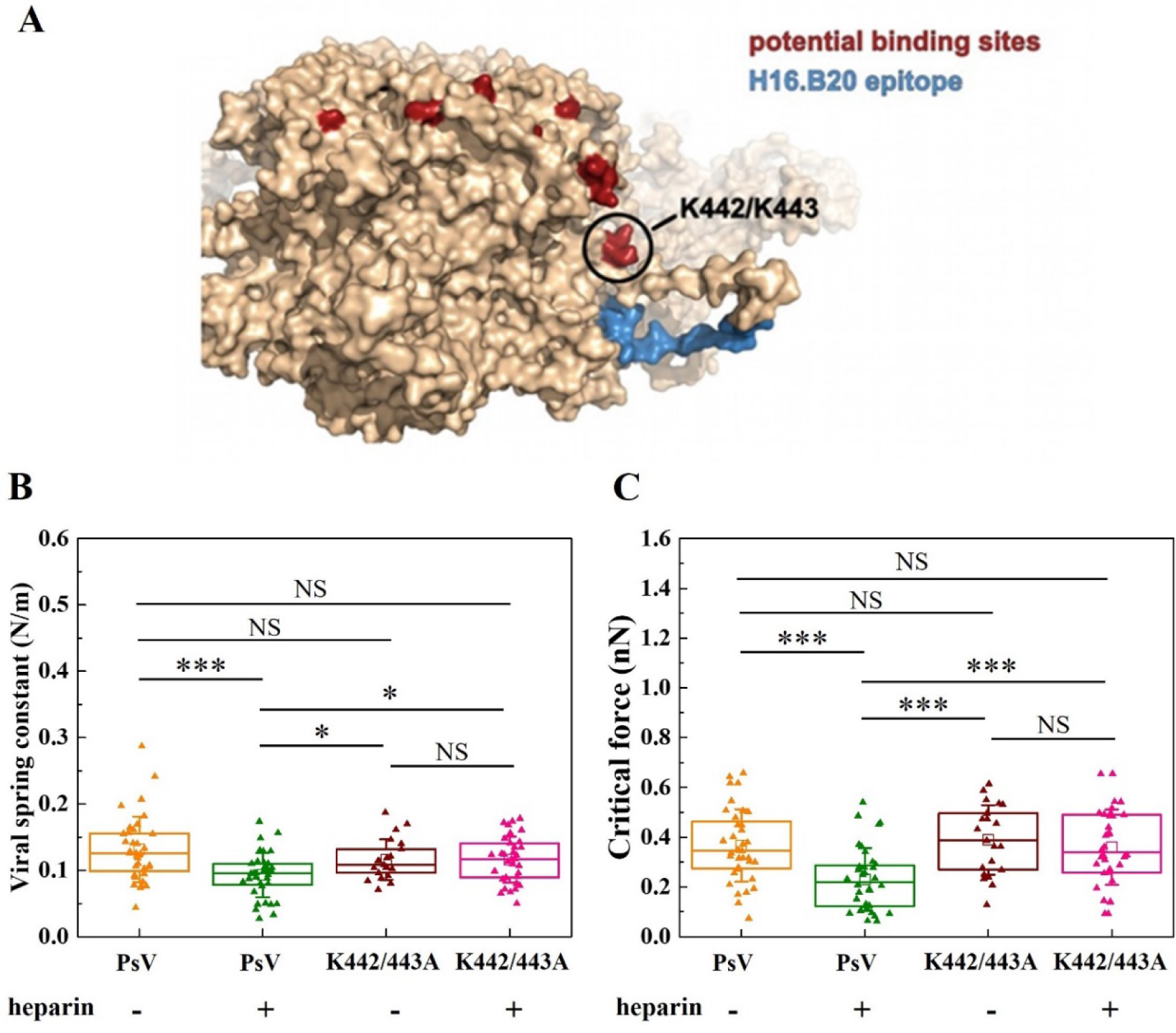
K442 and K443 are essential for activation of HPV16 through heparin. (A) Visualization of mutated site K442/443 on the L1 pentamer (5KEP) ^13^. Putative binding sites are in red ^24^ and the epitope of H16.B20 antibody is in blue ^48^. (B) Box plot of the viral spring constant of the HPV PsV and K442/443A PsV incubated with or without heparin. (C) Box plot of the critical force of the PsV and K442/443A PsV incubated with or without heparin. P values were determined by one-way ANOVA test. For the pairs showing statistically significant differences in viral spring constant, their P values for the comparison between PsV (N = 36 data points) and PsV with heparin (N = 36 data points), PsV with heparin and K442/443A (N = 21 data points), and PsV with heparin and K442/443A with heparin (N = 34 data points) are 3.59 × 10^−5^, 2.94 × 10^−2^, and 1.26 × 10^−2^, respectively. P values for the comparison of critical force between the corresponding pairs are 7.55 × 10^−5^, 7.42 × 10^−5^, and 1.89 × 10^−4^, respectively. P values are indicated by asterisks: p<0.001 (***), p<0.01 (**), p<0.05 (*), nonsignificant (NS).

According to this model, removing the K442/443 binding site would result in decreased or absent structural activation. Hence, we generated K442/443A mutant HPV16 PsV (HPV16 K442/443A). While these particles were similar to the WT in terms of particle morphology (Figure S3C), viral genome encapsidation (Figure S3B), and viral mechanical properties (Figure S2D-E), they were unable to infect cells (Figure S3A). Loss of infection was mostly unrelated to cell binding, which was only slightly reduced compared to the WT, as expected (Figure S3D-E) ^24, 25^. To examine, if the mutant would still be structurally activated upon GAG engagement, we tested whether HPV16 K442/443A VLPs exhibited a decrease of viral spring constant and critical force after heparin incubation. Importantly, this was not the case (Figure 4B-C) indicating that HPV16 K442/443A was not structurally activated, despite its ability to engage HS on cells (compare Figure S3D-E). Thus, most likely engagement of multiple HS binding sites on the capsid, including K442 and K443, is required for structural activation in line with our tension-force model.

### HDX-MS reveals heparin-induced stabilization in the intercapsomer canyon

To better understand the structural consequences induced in HPV16 capsids by heparin binding, HDX-MS was used to probe local conformational dynamics of PsV. HDX-MS usually compares different states of a protein, e.g. with and without ligand, through monitoring spontaneous exchange of protein backbone hydrogens for deuterium in a D_2_O-based buffer. This is observed by the mass increase of peptides, which are generated after the labelling reaction. The process probes the solvent accessibility as well as the involvement of amide hydrogens in hydrogen bonding and secondary structures. Changes in structural dynamics or engagement of amide hydrogens in ligand binding are hence reflected in increased (exposure / dynamics) or decreased (protection / stabilization) HDX^43^. For this, PsV were incubated in a D_2_O-based buffer alone and in the presence of excess heparin (1 mg/ml), which induces full structural activation.

HDX-MS detected multiple L1 peptides of distinct protein regions with decreased deuteration upon heparin engagement (Figure 5). The relatively small deuteration changes were however statistically significant and consistent across a number of peptides spanning the same protein region, thus conferring high confidence. Overall, most of the affected regions cluster towards both L1 N- and C-terminal regions (Figure 5A). These are known to sample multiple conformations and form the base of the canyon linking adjacent capsomers^49^. Indeed, the maximal deuteration differences (Figure S4) superimpose predominantly with the intercapsomere canyon of HPV16 in a structural model (pdb:7kzf)^49^ (Figure 5B-C). The pentameric core of the capsomer remained mostly unaffected, which fits well its description as a largely rigid structure in a flexible environment of the pseudosymmetrical capsid^49^. Importantly, one of the regions most perturbed by the binding of heparin was in fact the flexible C-terminal arm protruding into surrounding capsomers, the base of which contains the K442 and K443 residues involved in structural activation by heparin (Figure 5C, red mark).

**Figure 5:**
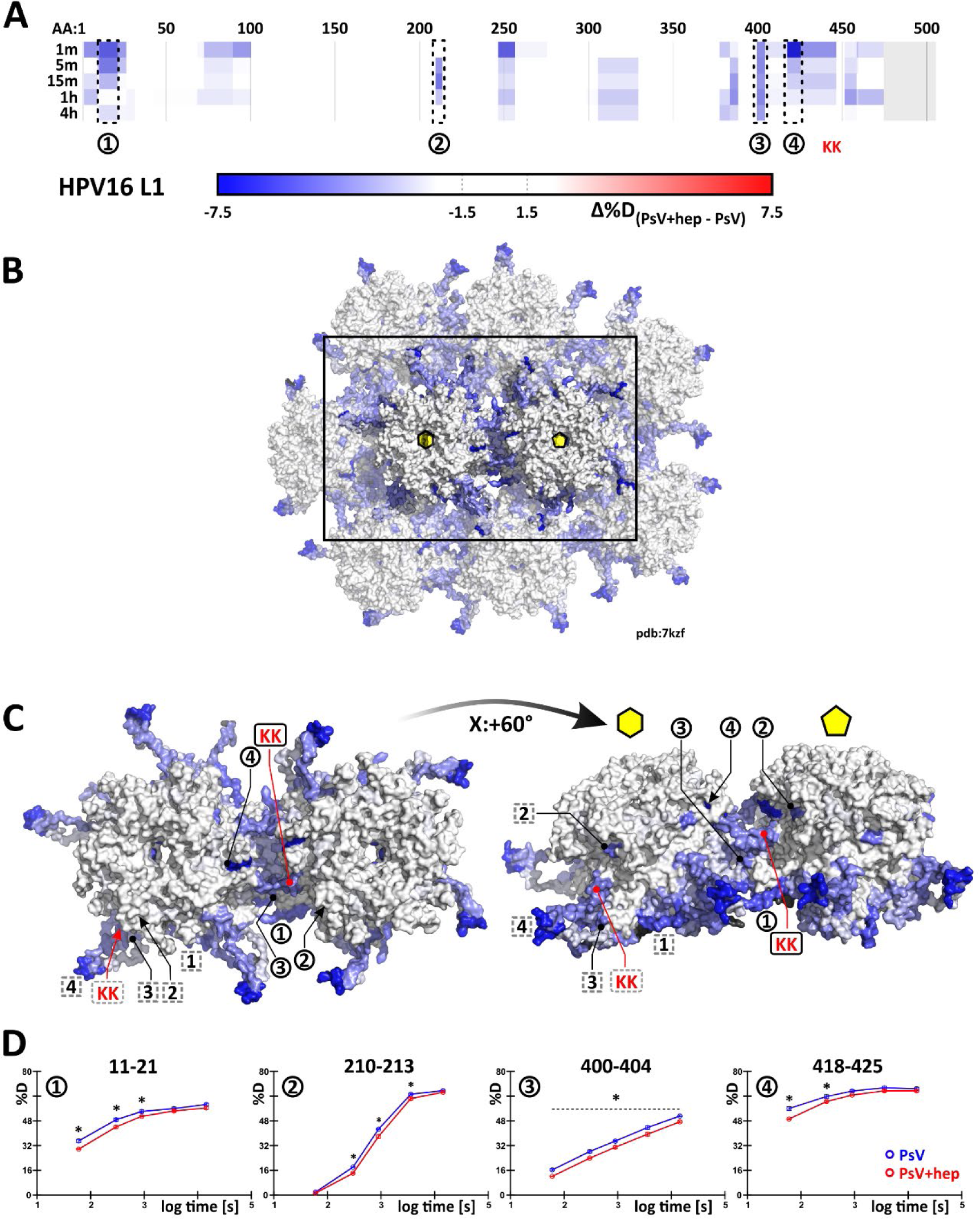
HDX-MS shows lowered deuteration inside the intercapsomer groove in the presence of heparin. (A) A subset of peptides describing all the observed changes in the course of multipoint HDX-MS experiment shown along the protein sequence, blue regions display decreased deuteration in the presence of heparin, grey area denotes missing data. Maximal observed differences are visualized symmetrically on the cryo-EM HPV16 structure. Hexavalent and pentavalent capsomers showed in broader context (B) and isolated (C). Regions whose example deuteration uptake plots are shown in (D) are marked on the asymmetric unit’s chain A (forming the pentavalent capsomer) – black circles and on the asymmetric unit’s chain C (part of the hexavalent capsomer), which is rotated about Z:+144° - grey dashed squares. Black regions on the structure denote missing data in B and C. Red marks the location of the K442/K443 patch, asterisk denotes statistical significance as described in the text.

In principle, the decrease of L1 backbone deuteration might at least partially be attributed to lowered solvent accessibility caused by an obstructing heparin molecule positioned in the capsid canyon as modelled in previous cryo-EM data^13^. However, electrostatic heparin/HPV interaction are mediated by basic amino acid side chains instead of the peptide backbone^24^. Such side-chain mediated interactions are often invisible in HDX-MS^50, 51^. In fact, functionally verified HS binding sites on the rim and at the side of L1 capsomers failed to exhibit a notable deuteration decrease (compare Figure 4A, Figure 5C, ^24^). Hence, the lowered deuteration more likely reflects decreased flexibility of L1 regions forming the canyon and corresponding rearrangement into more stable hydrogen bond networks upon structural activation.

In addition to L1, changes in the capsid-lumenal L2 would be plausible, since the L2 N-terminus must become exposed on the capsid surface for proteolytic processing and subsequent receptor switching^28, 31, 52^. Previous structural studies failed to conclusively identify L2 within the capsid probably due to its variable occupancy and suspected flexibility^12, 13, 49^. In our HDX-MS experiments, certain changes also on the lumenal side of L1 capsomers were observed, i.e. a decrease of deuteration upon the exposure to heparin both in the L1 amino acid stretch 306-327, next to which a more stable and thus better resolved density of a short stretch of putative L2 was previously identified in cryo-EM^49^, and in the region 247-256, which forms the inner lining of the capsomer pore (regions A and B in Figure S5A, respectively). In line with changes on the lumenal capsid side, HDX-MS data exhibited a decrease of deuteration in several regions of L2, likely due to lowered flexibility of the protein upon heparin engagement thus indicating that allosteric conformational changes indeed propagate to the L2 protein inside the capsid (Figure S5B). Notably, many regions without deuteration differences in L2 showed maximum deuteration within the first minute after labelling initiation (Figure S5C). This unequivocally confirmed a previously hypothesized very high flexibility and a degree of inherent structural disorder in these parts of L2^49^.

In summary, our HDX-MS data on the decreased exchange in L1 and L2 supports the relevance of the capsomer canyon and invading C-terminal arm in heparin binding and structural activation. Importantly, this data is consistent with the model of flexible conformations of the L1 C-terminal arm^49^, in which stabilization of extended conformations by e.g. heparin engagement would result in a notable size increase of the capsid. This would stabilise a more structured form with stronger hydrogen bonding, hence resulting in the decrease in deuterium exchange observed.

## Discussion

In this work, we probed the structural activation of HPV16 virions for infectious entry upon HS engagement from different angles: Structural activation of virus particles for functional receptor switching and entry upon interaction with heparin correlated with particles becoming significantly softer. Both required the engagement of HS with longer polysaccharide chains, and thus likely an engagement of several sites in the virus capsid by HS. As structural consequence of HS engagement, primarily a stabilization of the invading C-terminus of L1 into neighbouring pentamers was observed, likely in a more extended conformation, as this could account for softening of the particle. Thus, we propose that the strain introduced by the binding of stiff HS provides the force needed to reach and stabilize an extended conformation of the C-terminal arm of L1 linking capsomers, which in turn would facilitate subsequent alterations of the capsid and eventually receptor-switching and uptake (Figure 6). Counter intuitively, the HS-induced stabilization leads to a decrease in capsid stiffness. However, this stiffness decrease can be explained by the increase in size of the capsid, resulting in a more deformable particle.

**Figure 6:**
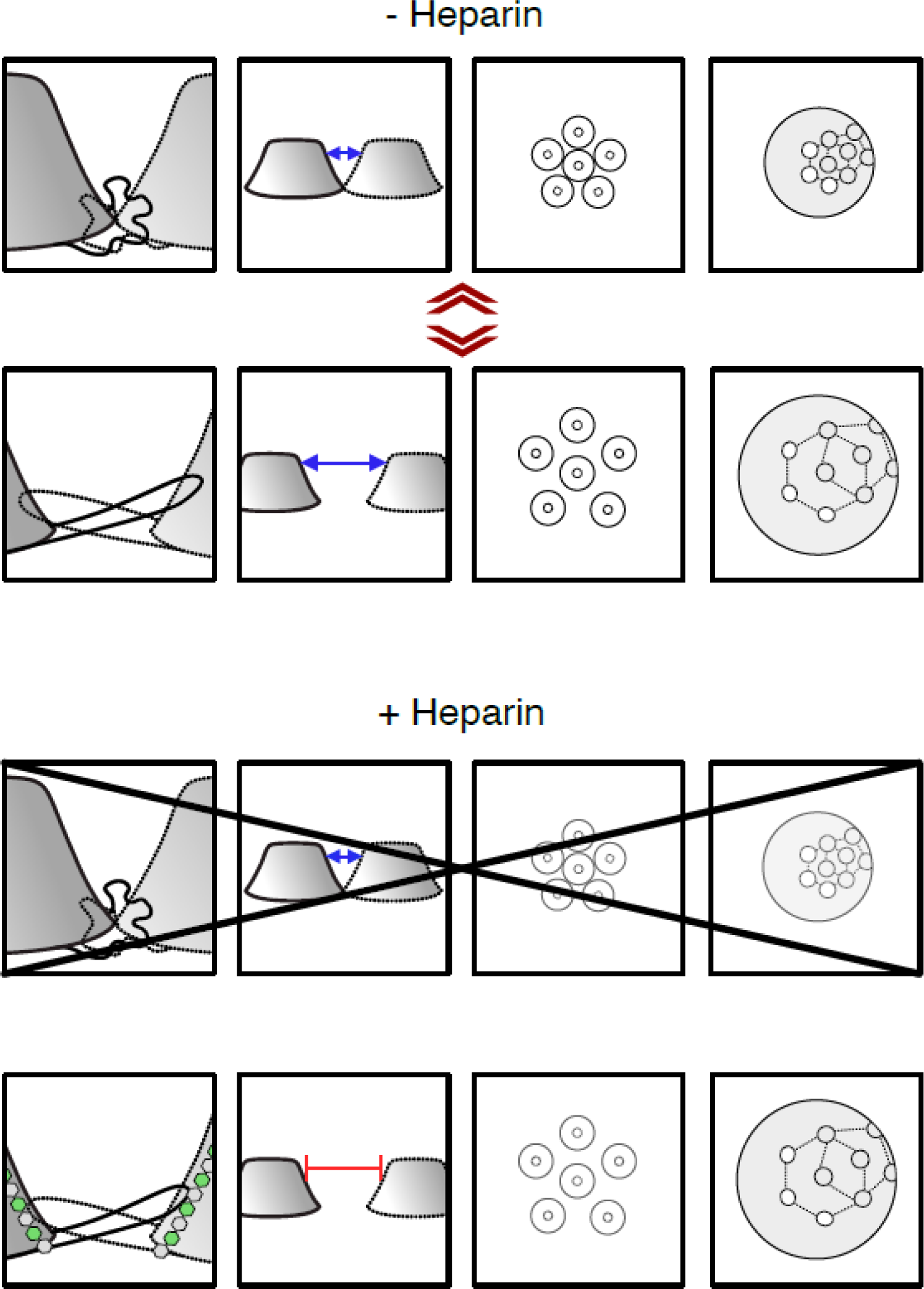
Model of HPV16 structural activation through HS/heparin. (A) Schematic of HPV16 pentamers. The pentamers are viewed from the side in column one and two and from the top in three and four. Column one displays the invading arm that forms a disulfide bond in the adjacent pentamer to stabilize the particle. The invading arm is very flexible, which constantly causes the pentamers to move towards and apart from each other (compare top and bottom). The so-called ‘particle breathing’. (B) Binding of a long HS/heparin molecule to several aligned binding sites including K442/443 stabilizes the invading arm in the most extended conformation, which reveals the H16.B20 epitope as well as the KLK8 cleavage site. The maximum interpentameric distance leads to an increased particle size and a softening in AFM, while additional hydrogen bonds in the stretched arms decrease HDX in this region.

Combining AFM nanoindentation, HDX-MS, and seedover assays, we demonstrated that structural activation by HS engagement is reversible and depends, both, on the length of the glycan and lysines 442/443 of the HPV16 L1 leading to stabilization of the intercapsomer canyon formed by the L1 invading C-terminus with lysines 442/443 at its base. While most protein-HS interactions require glycan chains as short as dp 4, and while short heparin fragments can bind to the HPV16 capsid^26, 47^, short heparin fragments were insufficient for HPV16 structural activation. Together, this implies that engagement of several binding sites by one HS molecule is crucial to induce or stabilize a particular conformation in the HPV16 capsid. No structural data on the engagement of long HS molecules to HPV16 capsids exists. Existing more indirect data including this study allows to infer whether binding of one HS molecule occurs to several distinct HS binding sites within one L1 molecule or to binding sites spanning different L1 molecules: As it provides means to increase affinity/avidity for interactions, multivalent HS binding is not uncommon for viruses^53, 54, 55^. Theoretically, this might explain the perceived lack of structural activation for HPV16, because low avidity binding of HS to capsids could result in low structural activation. However, since binding of HPV16 K442/443A to cellular HSPGs was efficient (this study and ^24, 25^), this appears unlikely. Moreover, since structural activation required engagement of K442/443, it is more plausible that one HS molecule binds simultaneously to several if not all binding sites in one L1 molecule in line with a previously proposed sequential engagement of binding sites^25^. Thus, we propose that a long stiff heparin/HS molecule binds to multiple binding sites on one L1 molecule, during which the heparin/HS molecule is bent and thus exerts forces on capsids. Here, the lowest binding loop, the lysines 442 and 443 would act as a handle for the GAG.

The residues 442 and 443 are directly adjacent to the invading arm of L1, which is crucial for particle stability and integrity by forming a covalently connected interpentameric network^15, 56, 57, 58^. Indeed, HDX-MS data indicated a general decrease in deuterium exchange of this region. Since the invading arm is a rather flexible structure that can exist in several different conformations^49^, HS engagement of K442/443 may stabilize one of those L1 C-terminal conformations in line with decreased deuteration. The different C-terminal arm conformations have been attributed to a phenomenon termed ‘particle breathing’, during which the L1 pentamers converge and diverge by about 7-9 Å^49^. If HS engagement would stabilize one of those L1 conformations, it would likely result in an observable change in the average diameter of the particles. Indeed, average particle size increased significantly upon engagement of long heparin polysaccharides but not of smaller oligomers (Figure S1A) ^13^, strongly suggesting that an extended conformation increasing the interpentameric distance was stabilized. An extended conformation of the L1 intercapsomer connection would also fit well with a softening of the virus particle, which was observed as reduced critical force and spring constant in AFM (Figure 2D-E). Moreover, the increased particle size is also in line with the enhanced accessibility of the HPV16.B20 epitope^26^.

Overall, our mechanistic model for structural activation of HPV16 particles fits well within the paradigm of viral entry, in which crucial epitopes of viral structural proteins are exposed upon cell interactions in a timely and spatially exact manner to facilitate the next of many steps in the entry program^59^. More specifically, our model may indeed explain how the immediate next steps in the entry program would be facilitated: Subsequent to binding to HSPGs, L1 is cleaved by kallikrein proteases, a capsid-lumenal N-terminal L2 peptide is exposed to the virion surface by the help of cyclophilins, which is eventually cleaved by secreted cellular furin^27, 28, 31^. A stabilized L1 C-terminus within an enlarged particle would increase the efficacy of the crucial kallikrein-8-facilitated proteolysis of L1 for cell entry^26, 27^. In turn, both, the enlarged capsomer distance and kallikrein cleavage of L1 would allow easier access of cyclophilins to the capsid lumen promoting the exposure of the L2 N-terminus to the capsid surface, which would allow efficient furin-mediated cleavage of L2 that is crucial for receptor switching. It is also plausible that allosteric transmission of structural changes within L1 to the capsid lumenal L2 would help cyclophilin-mediated exposure of the L2 N-terminus. In an elegant way, HPV16 has evolved to stabilize one of several alternating conformations in the capsid to promote efficient entry. Making use of these findings by designing small compound inhibitors of such ‘molecular gymnastics’ and adding those e.g. to lubricants may provide an additional strategy to reduce the burden of anogenital cancers.

## Methods

### Cell lines, plasmids, reagents and viruses

HeLa cells were from ATCC. HaCaT cells from N. Fusenig (DKFZ, Heidelberg, Germany) ^60^ and HEK293TT cells ^20^ were a kind gift from J. T. Schiller (NIH, Bethesda, USA). The plasmids pClneo-EGFP, p16L1h and p16Shell were kindly provided by C. Buck (NIH, Bethesda, USA) ^21, 61, 62, 63^. For maintenance, cells were kept in Dulbecco’s Modified Eagle’s Medium (DMEM, Thermo Fisher Scientific) which was complemented with 10 % fetal bovine serum (FBS, Biochrom). In case of HEK293TT cells the medium was supplemented with 400 µg/ml Hygromycin B. SYBR green master mix and BSA standard were purchased from Thermo Fisher Scientific. OptiMEM (Gibco), Lipofectamine 2000 (Invitrogen), Optical bottom 96w microplates (Greiner Bio-One).

### Virus preparation

HPV16 PsV were prepared according to ^21^. In short, p16Shell and pClneo-EGFP were transfected into HEK293TT cells. After 48 h, cells were harvested and lysed followed by maturation of the virus particles for 24 h. For purification, the particles were subjected to a linear OptiPrep (iodixanol, Sigma-Aldrich) gradient (25 – 39 % OptiPrep, 309600 x g, 5 h, SW60Ti rotor (Beckman Coulter). Purification of VLPs was realized accordingly, with the L1-only plasmid p16L1h but without addition of the reporter plasmid pClneo-EGFP. For the purposes of HDX-MS analysis, the particles were purified using a CsCl step gradient (27 % w/V and 38.8 % w/V CsCl in 10 mM Tris-HCl pH 7.4, 207570 x g, 3 h 50 min, 4 °C) followed by dialysis in Float-A-Lyzer devices (1 mL, Spectra/Por) against a total of 3 L HPV virion buffer (1x PBS, 635 mM NaCl, 0.9 mM CaCl_2_, 0.5 mM MgCl_2_, 2.1 mM KCl, pH 7.4), since iodixanol, the main component of OptiPrep, precipitates in the presence of acetonitrile used for peptide liquid chromatography (LC) separation.

### Glycosaminoglycans

If not indicated otherwise, heparin was from Sigma-Aldrich (H4784, mainly dp45 – dp51). To investigate the influence of chain length on structural activation a set of GAGs with a controlled length was utilized. Fondaparinux is a common clinically used heparin with dp5 (04191876, Viatris). The heparin oligomers dp14, dp18 and dp20 (HO14, HO18, HO20) and the HS fractions HS dp190 and dp40 (GAG-HS I and GAG-HS III) were from Iduron.

### Infection assays

About 4000 HeLa cells/well were seeded in 96-well optical bottom microplates 16 h prior to infection. To infect the cells, about 3 * 10^7^ HPV16 PsV were added and incubated at 37 °C for 2 h, before the medium was exchanged and replaced by DMEM (10 % FCS). Cells were fixed using 4 % paraformaldehyde (PFA, in PBS) 48 h p.i. Nuclei were stained with RedDot2 (VWR). Cells were imaged with the 10x objective of a Zeiss Axio Observer Z1 spinning disc microscope (Visitron Systems GmbH), which was equipped with a prime BSI camera (Photometrics) and a Yokogawa CSU22 spinning disc module. For each of three independent experiments 32 fields of view were imaged per condition. Infection was scored as GFP-positive cells/total cells in % in an automated manner using MATLAB-based InfectionCounter as previously reported ^64^. Experiments for inhibition of infection by different glycans was performed by preincubating HPV16 and the respective glycan at RT for 1 h, before infection of the cells was carried out as above.

### Structural activation assay (seedover)

Activation of HPV16 was tested as previously described ^26^. About 25000 HaCaT cells/well were seeded in 96-well optical bottom microplates 48 h prior to infection. Furthermore, 48 h before infection different HaCaT cells were supplemented with 50 mM NaClO_3_. This NaClO_3_ treated medium was renewed every day until the cells are fixed. On the day of infection, the cells in the 96-well plates were incubated with 20 mM EDTA (diluted in PBS) for 45 min at 37 °C. Afterwards, they were gently removed leaving the ECM behind. To test for their activation potential on HPV16, different glycans were incubated together with the virus for 1 h at RT. This mixture was added to the ECM and incubated for 1 h at 37 °C. About 4000 NaClO_3_ treated or control HaCaT cells were seeded in the 96-well plates to be infected by the ECM bound virus. 48 h p.i. the cells were fixed with 4 % PFA (in PBS). Microscopy-based scoring of infection was executed as described above. To investigate if the structural activation is reversible, the virus/glycan-mixture is incubated with 0.5 U heparinase I and III (H3917, Sigma-Aldrich) for 1 h at RT, before being added to the HaCaT ECM.

### AFM sample preparation

For glycan treatments, the heparin (H4784, Sigma-Aldrich) and Fondaparinux (04191876, Viatris) were solved at a concentration of 50 mg/ml in 10 mM HEPES buffer (pH 7.4). HPV16 particles were incubated with the glycan solution with 1 mg/ml final concentration for 1 hour at room temperature before performing AFM measurements. An equivalent volume of 10 mM HEPES to the glycan solution was added to an HPV16 solution as the untreated control group.

To enzymatically digest bound heparin, heparinase I and III Blend (H3917, Sigma-Aldrich) was solved with the concentration of 0.2 U/μl in the resuspension buffer, which was made of 20 mM Tris, 50 mM NaCl, 4 mM CaCl_2_, 0.01 % BSA and adjusted pH to 7.5. Subsequent to incubating HPV16 particles with heparin, 0.5 U of heparinase solution was added and incubated for another 1 hour at room temperature. An equivalent volume of PBS to the heparinase solution was added to a heparin-HPV16 solution as a control group.

### AFM imaging and nanoindentation

Prior to AFM measurement, different HPV16 particles stock solutions were first diluted to a concentration of 11.3 μg/ml. Generally, AFM imaging and nanoindentation experiments were conducted following the previous protocol^65^. Briefly, a droplet of 100 μL HPV solution was deposited on a hydrophobic glass coverslip and incubated for 15 min at room temperature before adding another 1 ml PBS to a liquid receptacle. Both AFM imaging and nanoindentation were performed in liquid at room temperature using an AFM (Nanowizard 4 model, JPK). Rectangular cantilevers (qp-BioAC CB2, Nanosensors) with nominal spring constant at 0.1 N/m and nominal tip radii below 10 nm were used and calibrated using the thermal noise method. Before nanoindentation, a series of AFM images were collected using quantitative imaging mode at setpoint 50 pN - 80 pN to centralize one isolated HPV particle and ensure the target particle was in proper shape. After completing a high-resolution image of the target particle, nanoindentation was performed on the clean substrate neighbouring the particle to check AFM tip cleanliness. Then, the AFM tip immediately pushed the centre of the target particle at a loading velocity of 300 nm/s until 3 nN force was reached. Finally, the particle was imaged again to inspect the structural state after the indentation. AFM images were processed using JPKSPM data processing software (Version 6.1.163, JPK) and corresponding force curves were processed by Origin software. Statistical analyses were performed using SPSS software (ver. 24.0, IBM).

### Affinity-based purification of HPV16

About 1.5 * 10^12^ of HPV16 PsV were subjected to affinity chromatography on HiTrap Heparin HP (1ml, VWR) using a NGC Quest System (BioRAD). While buffer A was composed of 10 mM sodium phosphate buffer pH 7.4, buffer B was additionally supplemented with 2 M NaCl. Sample application was performed in 15 % buffer B, followed by a wash-phase for 30 min (1 ml/min). Finally, the virus was eluted with a continuous 15 – 75 % buffer B gradient and collected in fractions. Virus fractions were pooled and used in the seedover assay.

### Virus labelling

HPV16 was fluorescently labelled as previously described ^66^. Briefly, HPV16 was incubated with AF568 succinimidylester (Invitrogen) in a 1:8 molar L1:dye ratio in the dark for 1 h at RT while rotating. Then, it was purified on an OptiPrep step gradient (15 %, 25 % and 39 %) at 225884 x *g* for 2 h at 16 °C. The labelled virus appeared as clear band and was collected. After addition of 4 % glycerol, it was snapfrozen in liquid nitrogen.

### Virus binding

One day prior to infection, 5000 HeLa cells were seeded on a coverslip. AF488-labeled HPV16 or HPV16_K442/443A was allowed to bind to the cells for 2 h. Afterwards, the cells were washed thrice with PBS and fixed in 4 % PFA. Cells were permeabilized with 0.1 % Triton X-100 for 15 min, followed by staining with phalloidin-AF647 (Sigma-Aldrich) in PHEM buffer for 30 min. Images were acquired as single slices with a spinning disc microscope (Visitron Systems GmbH) with the 40x objective. Particles were counted with the 3D object counter of ImageJ (version: 2.0.0-rc68/15.2e)^67^.

### VGE determination

Using 0.2 mg/ml proteinase K (Sigma-Aldrich) in HIRT buffer (400 mM NaCl, 10 mM Tris-HCl pH 7.4, 10 mM EDTA pH 8.0) DNA was released from HPV16 particles. To recover the DNA a Nucleo Spin Gel and PCR Clean up kit (Macherey-Nagel) was used and the DNA was eluted in 20 µl TE buffer (10 mM Tris, 1 mM EDTA, pH 7.5). PClneo-EGFP incorporation was investigated by quantitative PCR (ABI7500, Applied Biosystems) using a plasmid standard to determine the number of pseudogenomes. These were then related to particle amount of the input.

### HDX-MS analysis

PsV were pre-incubated for 1 h either with or without heparin (H4784, Sigma-Aldrich) at room temperature. To initiate deuterium labelling the samples were 6-fold diluted with the same buffer they were obtained in, only made of 99.9% D_2_O (150 mM NaCl, 4.8 mM KCl, 10 mM Na_2_HPO_4_, 1.8 mM KH_2_PO_4_, 0.9 mM CaCl_2_, 0.5 mM MgCl_2_, pD 7.2). This resulted in a final concentration of 0.5 µM L1 monomer in the form of PsV with or without 1 mg/ml heparin during deuterium labelling. The exchange reaction was left to proceed at room temperature until aliquots of 45 µl were removed at predetermined time points (1 min, 5 min, 15 min, 1 h and 4 h). In the aliquots, the exchange was immediately stopped by twofold dilution with ice-cold quench buffer (0.25 M glycine, 100 mM TCEP, 8 M urea, indicated pH 2.7), resulting in final pH 2.5. Importantly, heparin as well as DNA are highly negatively charged and thereby precipitate under HDX-MS quench conditions. Thus, we employed a strategy of Poliakov et al. ^68^, utilizing protamine sulphate as an additive in the HDX quench buffer and modified it for HDX-MS with online proteolytic digestion. Therefore, for samples with heparin, the quench buffer additionally contained 1 mg/ml protamine sulphate (P4020, Sigma-Aldrich). After 30 s incubation on ice, the samples were centrifuged at 10.000 x *g* for 1 min at 0 °C. Each supernatant was transferred to a fresh tube and flash frozen in liquid nitrogen. Low binding microtubes and low binding pipette tips (both Axygen) were used throughout for all handling of viral particles.

The frozen samples were quickly thawed and injected into a refrigerated (1°C) HPLC system (Agilent Infinity 1260, Agilent Technologies), through a porcine pepsin column (≥ 3200 units/mg, Sigma-Aldrich) in-house immobilized onto POROS-20AL perfusion resin (Thermo Scientific) as described previously ^69^, which was kept at 4°C. Pepsin digestion was performed at isocratic 200 µl/min flow rate (0.4 % formic acid in water). After the digestion, peptides were online desalted for 3 min on a peptide microtrap (OPTI-TRAP, Optimize Technologies) and then eluted on a reversed-phase analytical column (ZORBAX 300SB-C18, 0.5 x 35 mm, 3.5 µm, 300Å, Agilent Technologies). There LC separation proceeded at 25 µl/min flow rate through an 8 min gradient of 8– 30% solvent B, followed by a 3 min gradient of 30-90 % solvent B (solvent A: 0.4 % formic acid in water, solvent B: 0.4 % formic acid in acetonitrile). The outlet of the HPLC system was connected to an electrospray ionization (ESI) source of an Orbitrap Fusion Tribrid Mass Spectrometer (Thermo Scientific). The instrument was operated in positive ESI MS-only mode for deuterated samples, scan range 300-2000 *m/z*, using 4 microscans at resolving power setting 120,000. In a separate measurement on non-deuterated sample, the instrument was used in positive data-dependent ESI MS/MS mode with 30% HCD dissociation, 1 microscan and 240,000 resolving power setting for the identification of all peptides produced by non-specific pepsin cleavage.

In total 22 pmol and 50 pmol L1 protein were injected per MS and MS/MS analysis, respectively. To minimize sample carry-over on the protease column, two washing solutions were always injected between sample injections modified from Majumdar *et al.* ^70^ (wash solution 1: 5% acetonitrile, 5% isopropanol, 20% acetic acid; wash solution 2: 4 M Urea, 1 M glycine, pH 2.5). All HDX samples were analysed in technical triplicates, except for the 15 min time point for PsV without heparin, which was only measured in duplicate.

Peptides were identified from the MS/MS data by the Andromeda search algorithm implemented in MaxQuant (version 1.6.5.0) using a custom protein database containing the sequences of HPV16 L1 and L2 proteins. Deuterium uptake for the identified peptides was calculated with DeutEx (in-house developed), manually inspected and the statistical significance of the observed differences in deuteration was evaluated by applying an unpaired two-tailed Student’s T-test with single pooled variance evaluated with alpha ≤ 0.05 using the Holm-Šidák correction for multiple comparisons in Prism 8.0.1 (GraphPad Software). The processed data were visualized using MSTools (https://peterslab.org/MSTools/) ^71^ and open-source PyMol 2.6.0a0 (Schrödinger, Inc).

## Acknowledgments

CU and MS acknowledge funding through the Geman Research Foundation (DFG) within Research Group ‘ViroCarb’ (FOR2327): Glycans controlling non-enveloped virus infections’ (UE 183/1–1 &-2, SCHE 1552/3-2). MS acknowledges further funding through DFG within the Heisenberg-Program (SCHE 1552 6-1) and for research equipment (INST 211/1029-1). WHR and CU acknowledge funding through the EU ViruScan project and WHR also through the EU INFRAIA consortium MOSBRI. AK gratefully acknowledges postdoctoral research fellowship from the Alexander von Humboldt Foundation. The Leibniz Institute of Virology is supported by the Free and Hanseatic City Hamburg and the Federal Ministry of Health (Bundesministerium für Gesundheit, BMG). We are grateful to Hartmut Schlueter for access to the UKE proteomics core facility and use of the Orbitrap MS.

