## Supplemental Material for "Glycan-induced structural activation softens the human papillomavirus capsid for entry through reduction of intercapsomere flexibility"

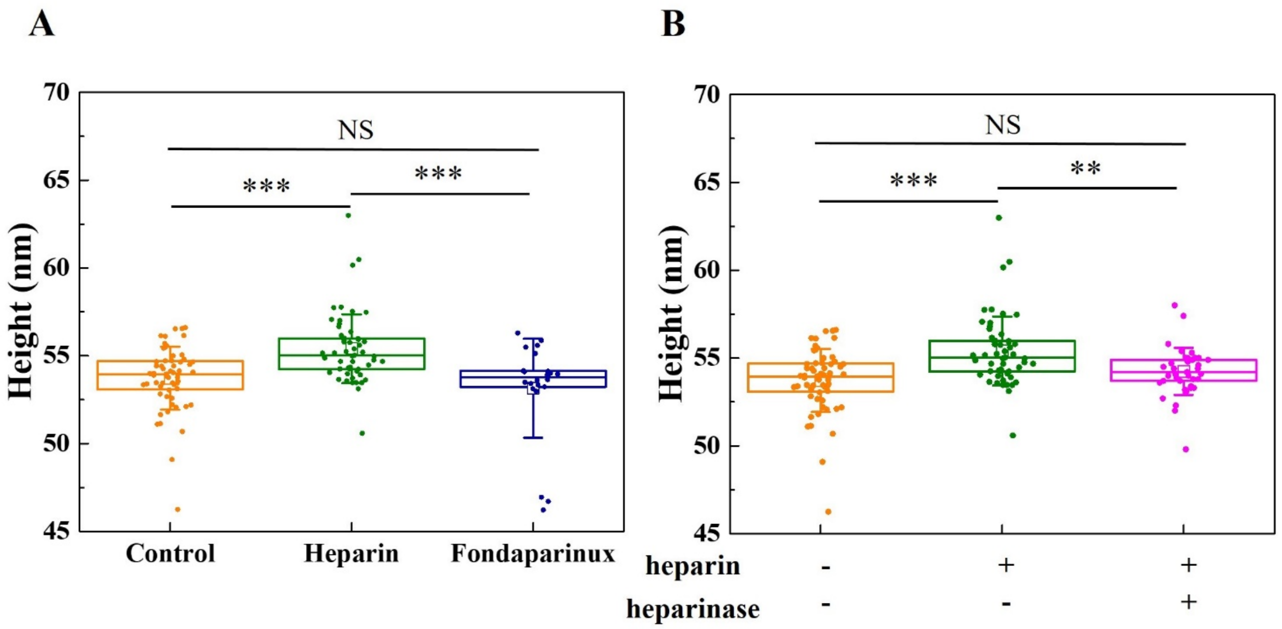


**Figure S1: Height of HPV VLPs after different GAGs treatments.**

(A) Comparison of the height of VLPs treated by different GAGs. (B) Comparison of the height of the VLPs treated by heparin and the VLPs treated by heparinase subsequent to heparin treatment. P values were determined by one-way ANOVA test. P values for the comparison of height between control and heparin, control and Fondaparinux, and heparin and Fondaparinux are $2.30 \times 10^{-5}$, 0.26, and $2.68 \times 10^{-5}$, respectively. P values for the comparison of height between control and heparin, control and heparinise subsequent to heparin, and heparin and heparinise subsequent to heparin are $8.33 \times 10^{-7}$, 0.14, and $1.64 \times 10^{-3}$ respectively. P values are indicated by asterisks: p<0.001 (***), p<0.01 (**), nonsignificant (NS).

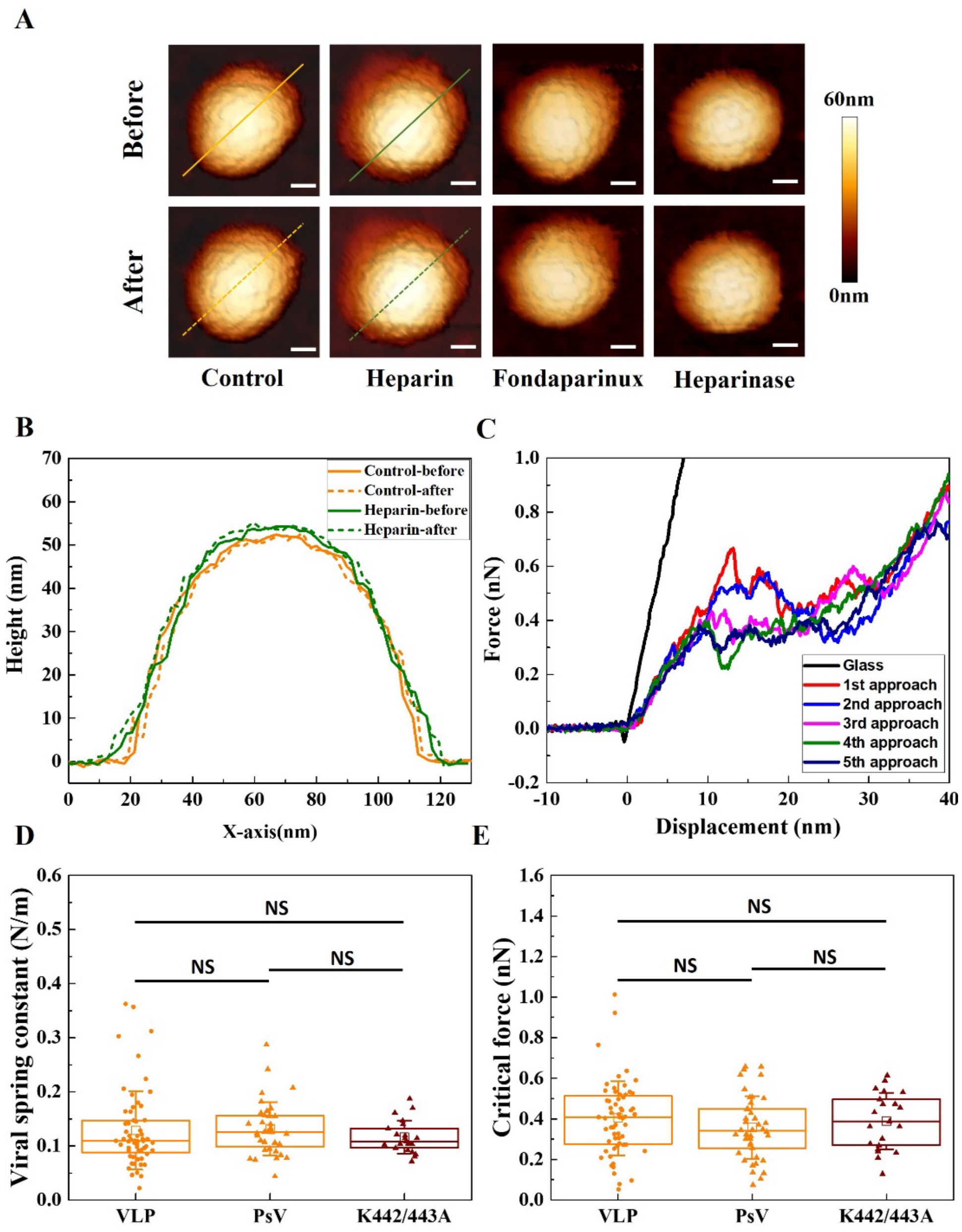


**Figure S2: Deformation of HPV particles under the applied force is elastic. HPV PsV and K442/443A PsV have similar mechanical properties as HPV VLP.**

(A) Representative AFM images of VLPs upon different GAG treatments before and after indentation. The scale bar is 20 nm. (B) Height profiles of untreated and heparin-treated VLPs before and after indentation. The height profiles of VLPs before and after indentation were acquired from the sections marked with solid and dashed lines in panel A). (C) Typical approach lines of the force-displacement curves of an untreated VLP for five successive indentations. (D) Box plot of the viral spring constant of VLP, PsV, K442/443A PsV. No significant differences in viral spring constant were found between the three groups by one-way ANOVA test. (E) Box plot of the critical force of VLP, PsV and K442/443A PsV. No significant differences in critical force were found between the three groups by one-way ANOVA test.

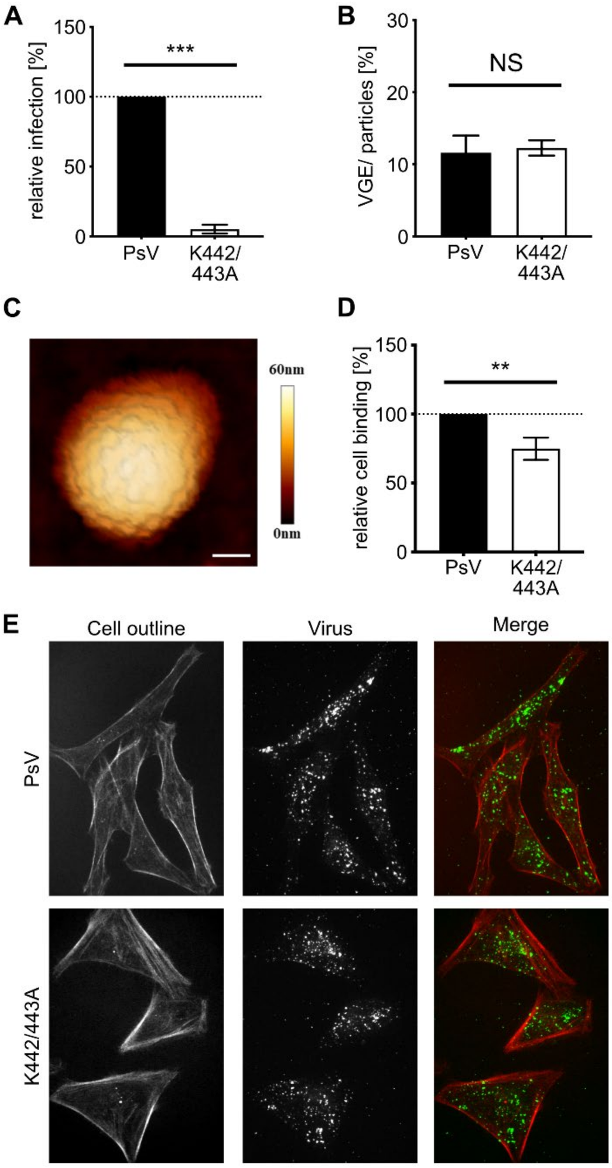


**Figure S3: Characterization of K442/443A PsV.**

(A) PsVs of either wt or mutant HPV16 were used to infect HeLa cells. GFP expression was detected 48 h p.i. by flow cytometry. Infected cells were displayed as average in % of total cells relative to the wt infection for three independent experiments ± SD. SD for the PsV control is 0 due to the normalization. (B) VGE were measured by qPCR using a viral pseudogenome standard. VGE values are displayed as average in % relative to the used particle amount for three independent experiments ± SD. (C) Representative AFM image of a K442/443A PsV particle. The scale bar is 20 nm. (D) Quantification of (E). 10 fields of view per replicate were analysed computationally to measure virus spots per cell area. Quantified spots were displayed as average in % relative to the wt for three independent experiments ± SD. SD for the PsV control is 0 due to the normalization. (E) PsV of either wt or mutant were fluorescently labeled and exposed to HeLa cells with the same amounts. The samples were analyzed with confocal microscopy 2 h post binding. Representative images were displayed as maximum intensity projections of confocal stacks. Cell outlines are derived from phalloidin staining. P values are indicated by asterisks: p<0.001 (***), p<0.01 (**), nonsignificant (NS).

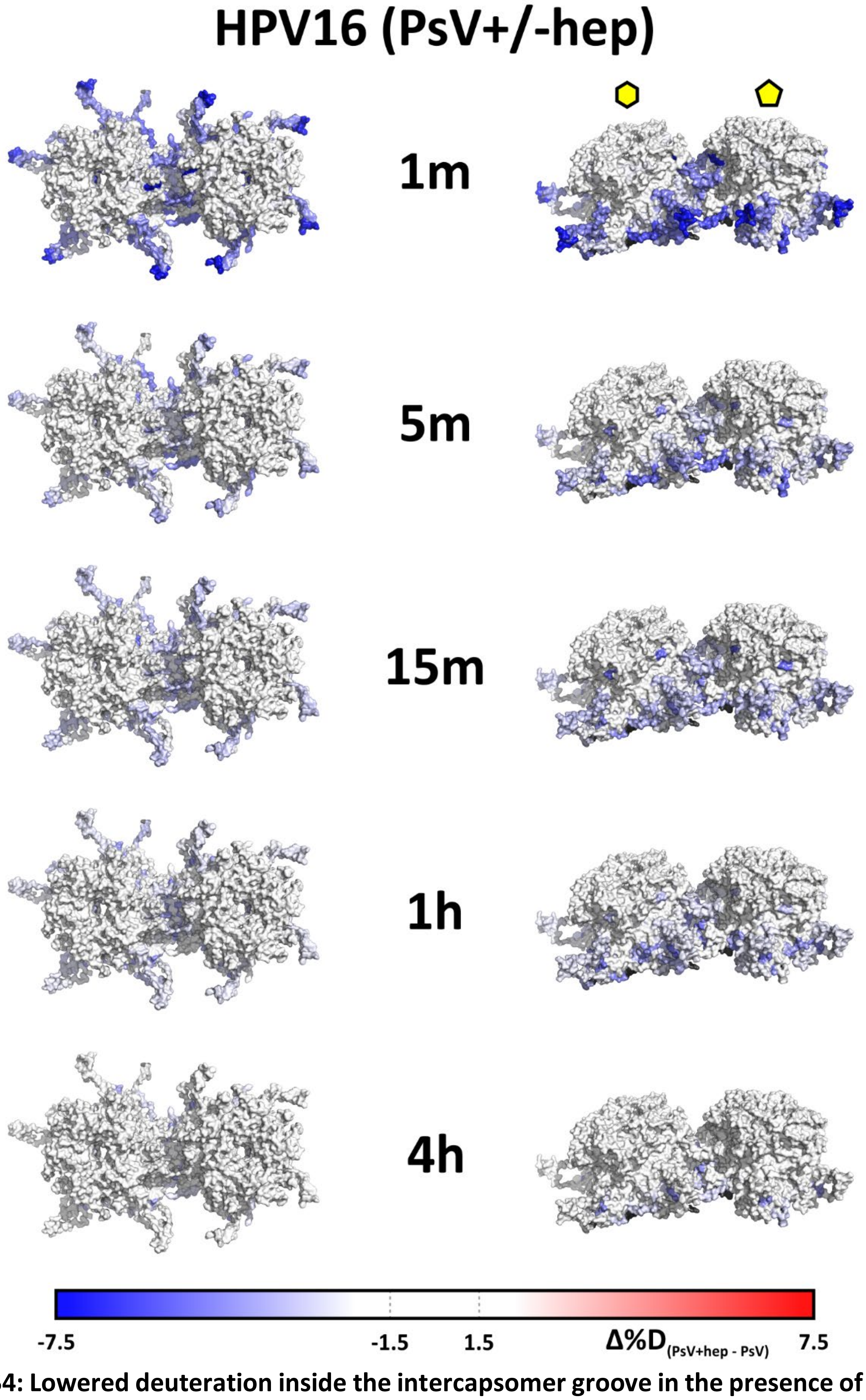


**Figure S4: Lowered deuteration inside the intercapsomer groove in the presence of heparin observed over the course of the HDX.**

Maximal observed differences in deuteration are visualized symmetrically on the cryo-EM HPV16 structure - hexavalent and pentavalent capsomers showed in isolation. Black regions on the structure denote missing data.

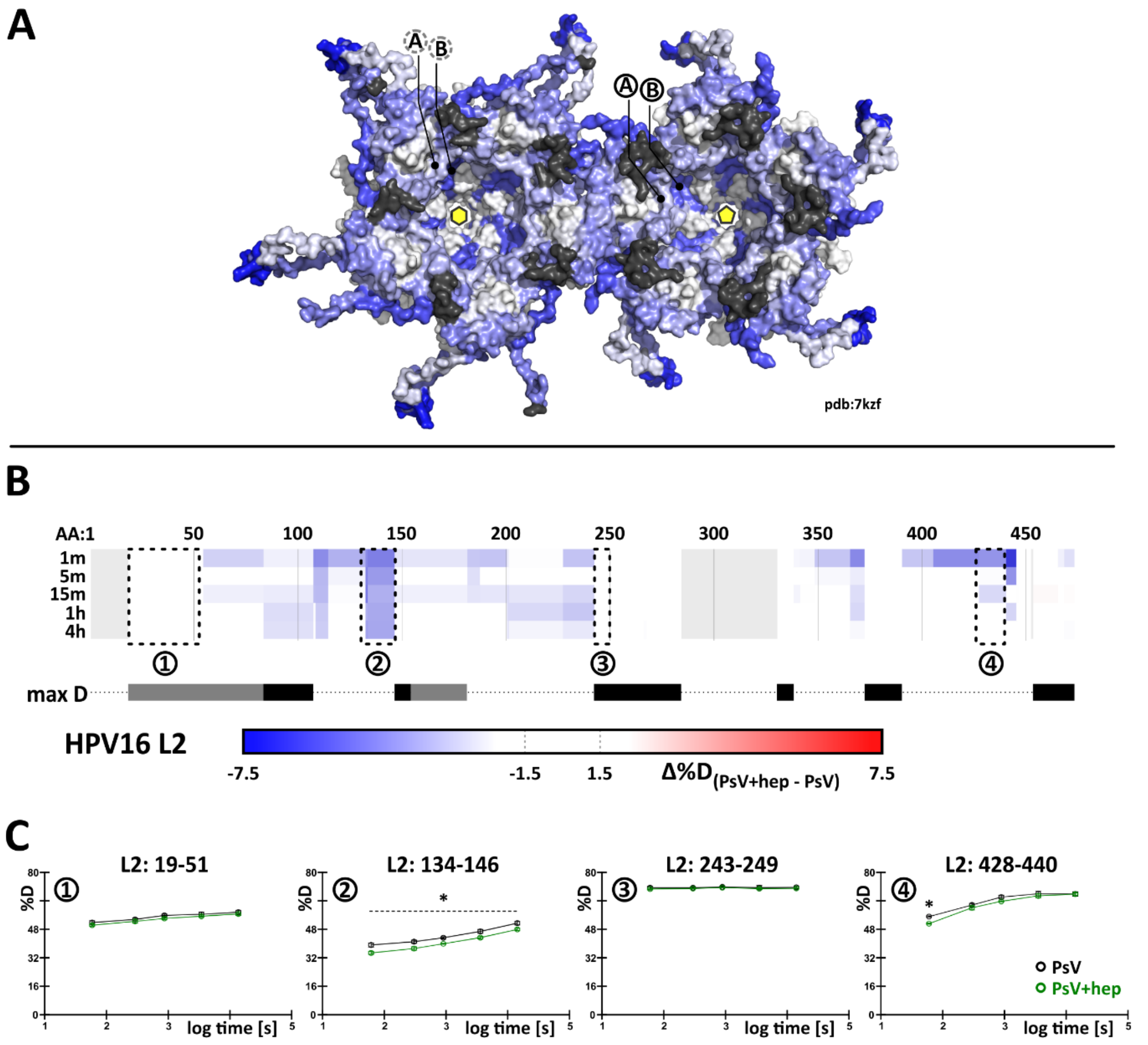


**Figure S5: HDX-MS observed lowered deuteration inside the capsid.**

(A) Decreased deuteration observed on the lumenal side of the capsid. Regions A and B show the region 306-327 and 247-256, respectively, marked on chain A (black circles) and chain C (dashed grey square) as described in Figure 5. Black regions mark areas with lacking data. (B,C) While missing high-resolution structural data, L2 protein also showed several regions with lowered deuteration, but additionally multiple regions with maximal deuteration achieved immediately (region 3, black bars) or almost immediately (region 1, grey bars) indicating unstructured / disordered regions. Light grey areas in (B) mark missing data.
